# PROCYCLIC *Trypanosoma brucei* CELL CYCLE is IMPAIRED at the G1 STAGE by the PRESENCE of POLY (ADP-RIBOSE) in the NUCLEUS

**DOI:** 10.1101/452730

**Authors:** Mariana Schlesinger, Salomé C. Vilchez Larrea, Silvia H. Fernández Villamil

## Abstract

Previously we demonstrated that an excess of poly (ADP-ribose) in the nucleus makes procyclic parasites more sensitive to hydrogen peroxide. However, the effect of an altered-PAR metabolism under standard conditions has not been addressed yet. Here we have analyzed the behavior of the growth curve of transgenic parasites that present this phenotype and studied cell cycle progression in synchronized cultures by flow cytometry and immunofluorescence. We have demonstrated that an excess of nuclear poly (ADP-ribose) produces a delay in the G1 phase of the cell cycle. Moreover, for the first time we have shown that poly (ADP-ribose) occurs at specific points very close to the mature basal body, suggesting there could be a link between the kinetoplast and poly (ADP-ribose) metabolism.

## 1 Introduction

*Trypanosoma brucei* is the etiological agent of the Human African trypanosomiasis, also known as sleeping sickness, and Nagana in animals. This parasitic disease is transmitted by the tsetse fly and affects mostly poor populations living in the Sub-Saharan Africa. According to the World Health Organization, 65 million people are at risk of infection. Its diagnosis is complex and this infection can lead to death when untreated.

Poly (ADP-ribose) polymerase (PARP) is responsible for the synthesis of polymers of (ADP-ribose) (PAR), which are involved in a wide range of cellular processes such as preservation of genome integrity, DNA damage signaling and repair, molecular switch between distinct cell death pathways and cell cycle progression. Only one *parp* gene is encoded within the *T. brucei*’s genome, called *Tb*PARP. In humans, however, there are 17 members of this protein family (*h*PARP). A sequence identity analysis of the catalytic domain evidenced that *Tb*PARP is similar to *h*PARP-1, *h*PARP-2 and *h*PARP-3 [1].

The enzyme in charge of degrading PAR and restoring the basal state of the cell is the Poly (ADP-ribose) glycohydrolase (PARG), which possesses an endo- and exo-glycohydrolase activity. The only member present in *T. brucei* is called *Tb*PARG. In humans, different isoforms derived from a single gene could be identified in the cytoplasm and the mitochondria.

In our previous study we have shown that *Tb*PARG occurs always in the nucleus, while *Tb*PARP localizes in the cytoplasm and migrates to the nucleus only after genomic damage [2]. In this organelle *Tb*PARP synthesizes PAR [3]. We have also analyzed the effect of hydrogen peroxide on procyclic cultures and demonstrated that a diminished *Tb*PARP activity conferred resistance to these parasites. In contrast, over-expression of this protein as well as silencing of *Tb*PARG expression lead to nuclear PAR accumulation, which resulted in an increase in their sensitivity. Even more, we have demonstrated that this genotoxic agent triggered a different death pathway in these transgenic lines when compared to the wild type parasites.

Our last work focused on the response to hydrogen peroxide of procyclic parasites with altered polymer levels. However, the effect of a modified PAR metabolism in standard conditions remained elusive. In the present work we have studied how nuclear PAR accumulation affects the trypanosomatid cell cycle progression. Moreover, we have shown for the first time that PAR synthesis occurs close to the kinetoplast and that *Tb*PARP colocalizes with the basal body in *T. brucei* procyclic parasites.

## 2 Materials and Methods

### 2.1 Parasite cultures

*T. brucei* procyclic strain 29–13 [4] was cultured at 28 °C in SDM-79 (Bioscience) containing 10 % (v/v) FCS and 0002 % hemin.

The transgenic lines p2216-TbPARP (which over-expresses the fusion protein *Tb*PARP-eYFP) and RNAi-TbPARG (with down-regulated TbPARG expression) previously obtained in our laboratory [2] were cultured in the same medium with the addition of 20 μg/mL zeocin.

Cell survival of transgenic cultures induced for 3 days with tetracycline (1.0 μg/mL) was determined counting motile parasites using a Neubauer chamber.

2.1.1-Synchronization, cell cycle and cell death analysis by Flow Cytometry.

Synchronized cultures were obtained as described by Chowdhury et. al [5]. A 10 mL culture (5×10^6^ cell/mL) was incubated in SDM-79 with 0.2 mM Hydroxiurea (HU) for 12 h at 28°C. After HU removal, parasites were washed twice and incubated in culture media for 12 h.

Propidium Iodide (PI) staining and Flow Cytometry: For fixation, cells (1 ml) were washed with PBS-EDTA (2 mM) and incubated in 70% cold ethanol for 24 h at 4°C. Then, cells were centrifuged at 15680 g for 10 min at 4°C and resuspended in PBS (1 ml) and stored at 4 °C. The same day of the analysis cells were harvested and stained with stain buffer solution (PBS 1X-2 mM EDTA, 200 μg/ml RNAsa, 50 μg/ml PI) for 30 min at 37 °C in the dark. Cell counting was carried out by flow cytometry (FACS Canto II, BD Pharmigen). Results were analyzed with Cyflogic v 1.2.1 software.

Cell death (apoptosis-like and necrosis pathways) of 3 day-induced cultures was assessed with FITC Annexin V Apoptosis detection kit from BD Pharmingen TM (San Jose, CA, USA), followed by flow cytometry analysis (FACS Aria, Becton Dickinson). Results were analyzed with Cyflogic v 1.2.1 software.

### 2.2 Immunofluorescence

Parasites were fixed with 3.8% (W/V) formaldehyde in PBS at 4°C, permeabilized with fresh PBS-0.1%Triton X-100 and blocked for 1 h at room temperature. PAR was detected with 1:500 primary anti-PAR antibody (BD) followed by 1:600 Alexa Fluor 594 goat anti-rabbit IgG antibody (Invitrogen). TbPARP was detected with 1:100 primary anti-TbPARP antibody (GenScript) followed by 1:600 Alexa Fluor 488 goat anti-rabbit IgG antibody (Invitrogen). Excess of antibody was removed by 3×5-min washes in PBS, and nuclear and kinetoplast DNA were stained with DAPI (2 μg/mL) (Sigma) in PBS. Coverslips were washed with distilled water and mounted in Mowiol and then visualized using an Olympus BX41 microscope.

Cytoskeleton extraction: Parasites were washed with PBS and attached to Poly-L-lysine (10mM) pre-treated slides. Incubation for 5 min with PEME buffer (0.1 M PIPES pH6.9; 1% P-40; 2 mM EGTA; 1 mM MgSO_4_ and 0.1 mM EDTA) was carried out for cytoskeleton extraction. After 3×5-min washes in PBS, cells were fixed with methanol for 30 min at 20 °C. A regular immunofluorescence protocol was followed as mentioned above from the blocking step after washing 2×5-min with PBS.

PAR was detected with 1:500 primary anti-PAR antibody (BD) and TbPARP was detected with 1:50 primary anti-TbPARP antibody (GenScript) followed by 1:600 Alexa Fluor 488 goat anti-rabbit IgG antibody (Invitrogen). The mature basal body was detected with 1:600 primary anti-α-Tyrosinated Tubulin antibody (YL 1/2) (Pierce Antibodies) followed by 1:250 Alexa Fluor 594 goat anti-rat IgG antibody (Invitrogen). Nuclear and kinetoplast DNA were stained with DAPI (2 μg/mL) (Sigma).

## 3 Results

### 3.1 Altered PAR metabolism causes a delay in procyclic cell cycle

Previously, we have obtained stable transgenic lines of procyclic *T. brucei* cultures which either over-express *Tb*PARP fused to eYFP (p2216-PARP) or down-regulate *Tb*PARG expression (RNAi-PARG). Both transgenic parasites have similar phenotypes, with alterations in PAR metabolism [2]. New studies carried out in the present work have shown that these parasites exhibit a delay in the growth curve (Fig. 1). The culture growth rate of the p2216-PARP induced parasites started diminishing as early as the second day post-induction, and at day three, the number of parasites in the uninduced cultures almost doubled the number of the induced ones (Fig 1A). This outcome was specific to *Tb*PARP over-expression, since when only eYFP was over-expressed (p2216-eYFP) the growth rate of the induced and uninduced cultures was similar. On the other hand, silencing of *Tb*PARG (RNAi-PARG) also caused a 50% decrease in the growth rate at day three of induction. This reduction in cell density at day three post induction could have two different causes: parasites with a modified PAR metabolism could either increase the cell death rate than PAR-unaltered microorganisms, or simply could grow up more slowly. We analyzed parasite motility as a sign of cell viability in three day cultures and no difference in the percentage of motile parasites was observed between the induced parasites and the uninduced control groups (Fig 1B). When we analyzed the presence of death markers by flow cytometry, the percentage of the dying populations (upper left, upper right and lower right quadrants) was in consonance with that obtained by estimation of cell viability (Fig 1C). This evidence supports the hypothesis that there is a delay in the cell cycle progression rather than an increase in the mortality rate.

**Figure 1.**
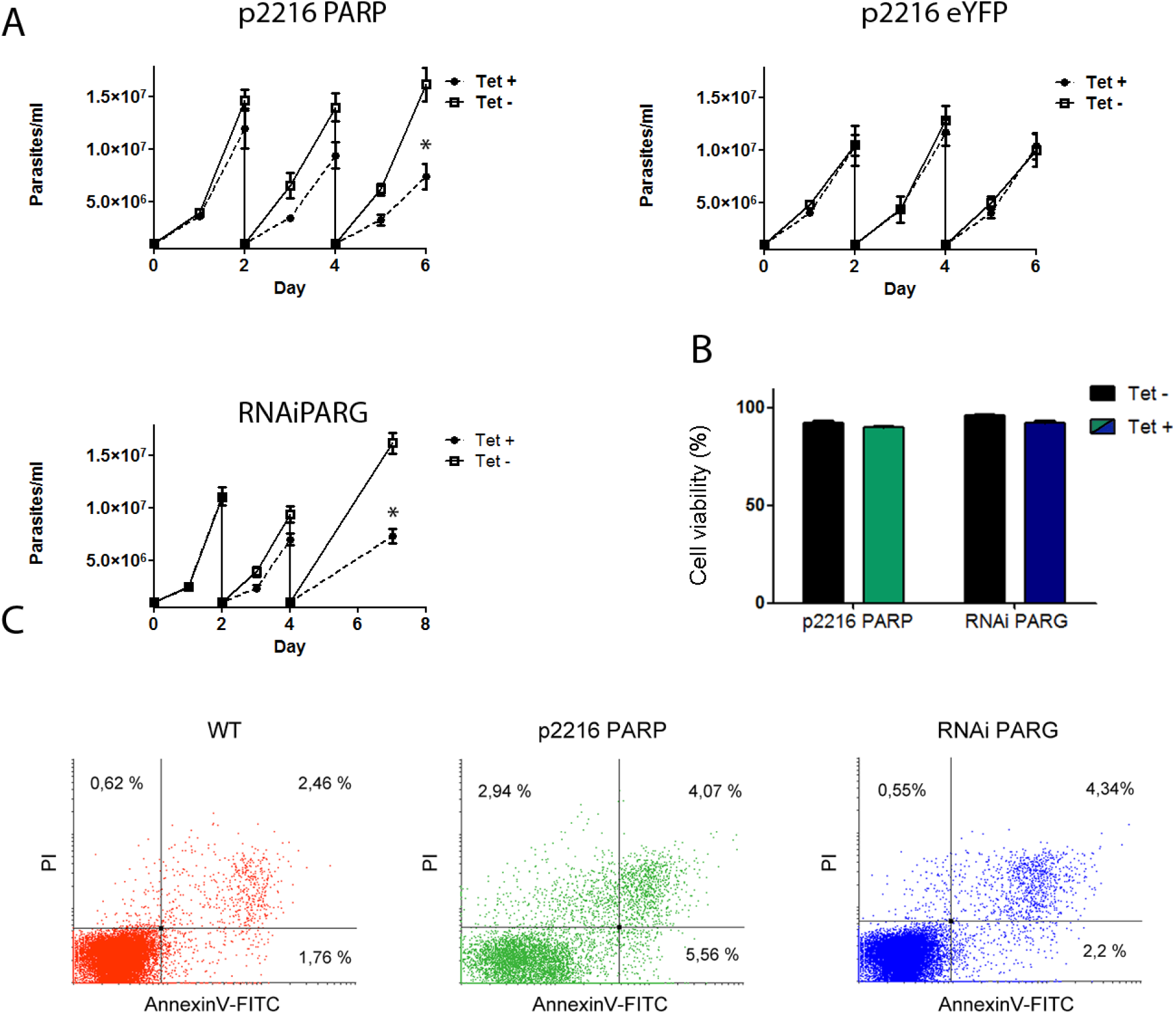
**Procyclic transgenic lines p2216-PARP and RNAi PARG**. (A) Representative growth curves of tetracycline-induced (Tet +) and uninduced (Tet -) parasites of p2216-PARP, p2216-eYFP and RNAi-PARG transgenic lines. Three independent experiments were carried out for each case (B) Cell viability was assessed by measuring the number of motile parasites respect to the total number in three day-induced cultures (Tet +) and uninduced controls (Tet -). Three independent experiments were carried out for each case. (C) Cell death analysis by flow cytometry of wild type and three-day induced transgenic cultures. Parasite populations were identified by staining with propidium iodide (PI) and Annexin V-FITC conjugate. The left quadrant of the p2216-PARP culture is moved to the right because of the intrinsic fluorescence from the PARP-eYFP fusion protein. Representative experiment of three independent assays.

With the aim to study whether the cell cycle is arrested at a particular phase in three-day induced parasites, we synchronized the cultures with hydroxyurea (HU) for 12 hours. Procyclic parasites are normally arrested at the end of phase S with HU, as opposed to other organisms which arrest at the boundary of the G1/S transition. It has been reported that once the HU is removed, procyclic parasite cultures remain synchronized for around 12 hours and the different phases can be identified by flow cytometry and Propidium iodide (PI) stain [5]. The approximate time period of each phase is schematized (Fig 2A).

**Figure 2.**
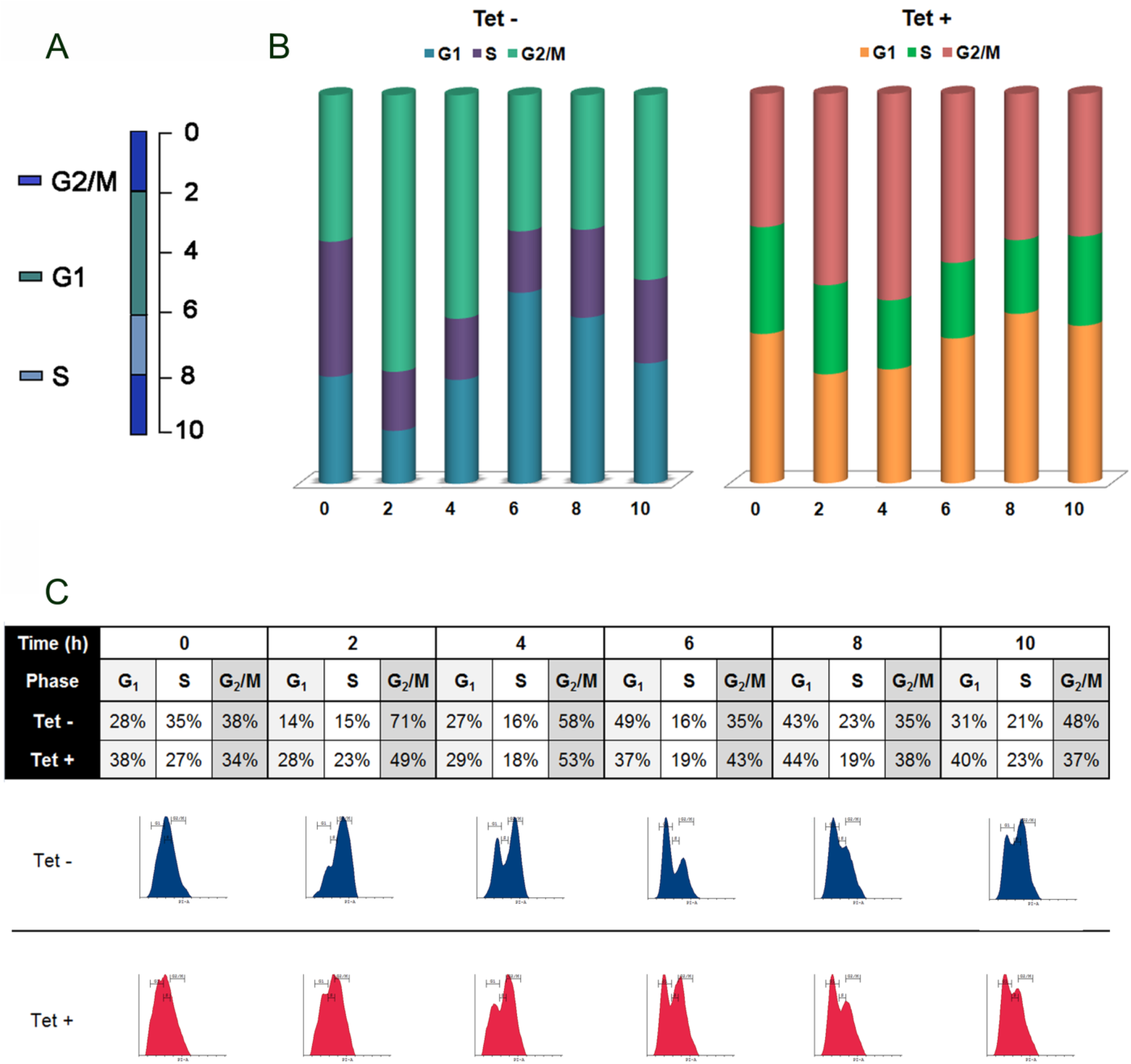
**Flow cytometry of p2216-PARP cultures synchronized with hydroxyurea (HU).** Flow cytometry of three day-induced (Tet +) and uninduced (Tet-) TbPARP over-expressing (p2216-PARP) cultures synchronized with 0.2 mM hydroxyurea (HU) for 12 hours. Samples were taken every two hours after HU removal to evaluate cell cycle progression. (A) Diagram showing the approximate time course of the cell cycle after HU removal. (B) Bar graph representing the percentage of the population in each cell cycle stage from the histograms at the times indicated. (C) Histograms obtained by flow cytometry based on the IP fluorescence signal. The table above displays the percentage of population for each phase (G1, S and G2/M). A representative experiment of three independent experiments is shown.

After HU was removed from synchronized p2216-PARP transgenic parasites, control cultures (uninduced) went through the S phase and entered the G2/M phase within the first 2 hours (Fig 2B and C). In the next hours (from 2 to 6) the G1 phase population kept increasing as a new cycle initiated. After that, the proportion of parasites in phase S began to augment (8 h) and within the last hours the culture progressively lost its synchrony while it proceeded through phase G2/M. Even though synchronized p2216-PARP transgenic cultures induced for three days presented the same pattern as the control groups, the cell cycle progression was delayed with an evident accumulation of the G1 population at hours 0, 2 and 10, as compared to uninduced parasites (Fig 2B and C).

Synchronized non induced RNAi-PARG cultures presented a cell cycle progression similar to that observed for the control group of p2216-PARP cultures (Fig 3A and B). Within the first 2 hours after HU removal the population in phase G2/M augmented. After that, G1 population started increasing until hour 6. At hour 7 the number of parasites in phases S and G2/M augmented considerably, something that could not be evidenced in the prior experiment with the p2216-PARP cultures. Finally the G1 population started increasing once again. Synchronized three day-induced RNAi-PARG cultures showed a delay in the transition from G1 to S phase, resulting in an increment of the G1 population immediately after the arrest was over (time 0), and at hours 7 and 8. Besides these observations, no apparent difference was observed between induced and uninduced cultures until hour 6, while the G1 population of the control culture kept increasing. Altogether, both experiments evidenced that synchronized parasites with a disrupted PAR metabolism have difficulties in the progression through the G1 phase, a phenomenon that is accentuated at time 0 probably due to the effect of the HU.

**Figure 3.**
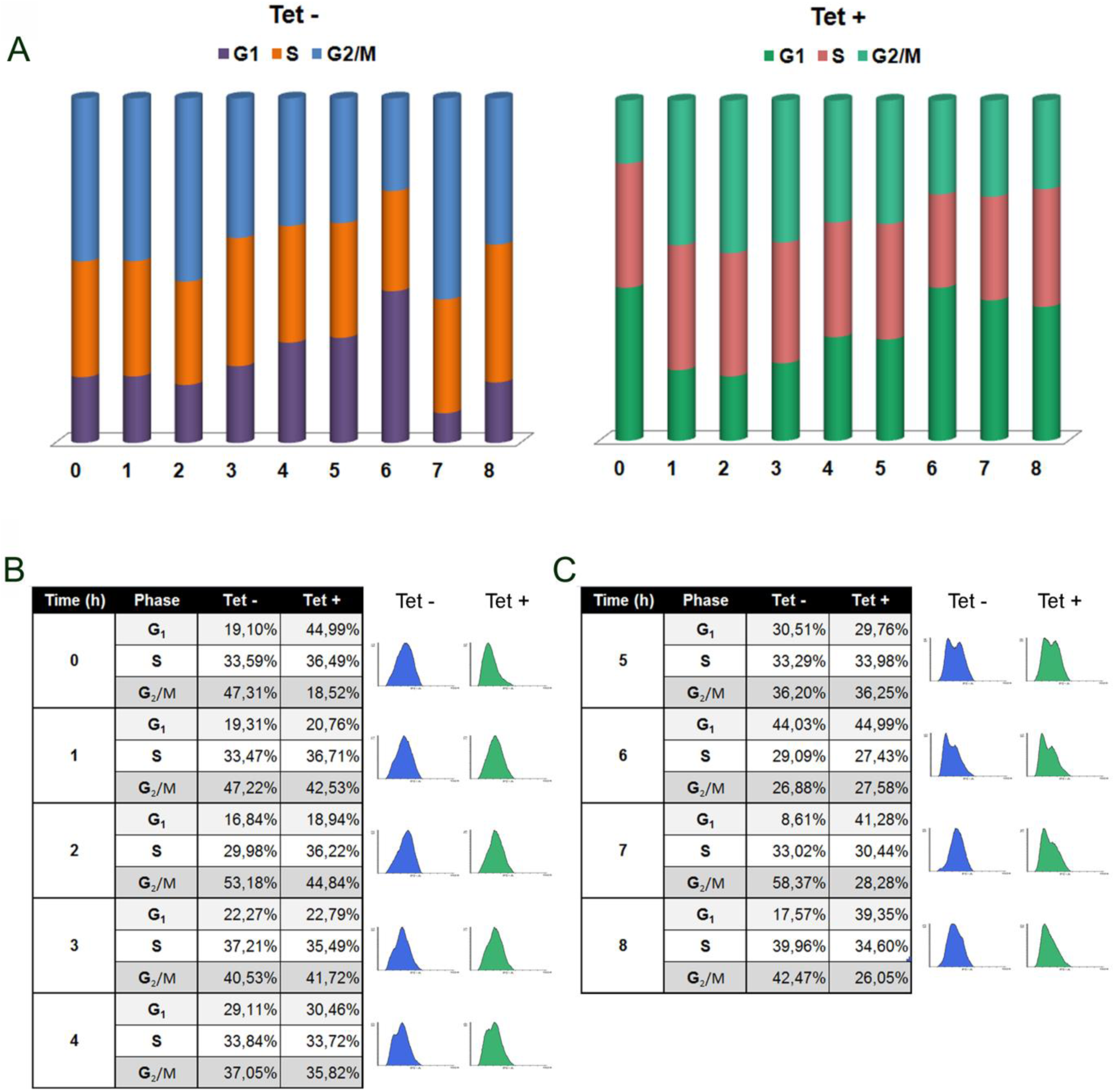
**Flow cytometry of RNAiPARG cultures synchronized with hydroxyurea (HU).** Flow cytometry of three day-induced (Tet +) and uninduced (Tet-) *Tb*PARG down-regulated (RNAiPARG) cultures synchronized with 0.2 mM hydroxyurea (HU) for 12 hours. Samples were taken every hour after HU removal to evaluate cell cycle progression. (A) Bar graph representing the percentage of the population in each phase of the cell cycle at the times indicated. (B) Histograms obtained by flow cytometry based on the IP fluorescence signal. The table displays the percentage of population for each phase (G1, S and G2/M). A representative experiment of three independent experiments is shown.

### 3.2 Extra nuclear PAR is synthesized near the kinetoplast

In our previous studies, we showed that *Tb*PARP is localized in the cytoplasm under basal conditions, usually in a region between the nucleus and the kinetoplast. *Tb*PARG, on the other hand, resides always in the nucleus [2]. We have also noticed that PAR cytoplasmic localization presents particularly one mark close to the kinetoplast, something that was confirmed here by IFI (Fig 4 A). Strikingly, cytoskeleton extractions from wild type procyclic parasites displayed polymer signal in two points next to the mature basal bodies (Fig 4B). The commercial monoclonal anti-YL1/2 antibody specifically detects tyrosinated α-tubuline, allowing the recognition of the posterior end of the parasite cell as well as the mature basal body fibers [6, 7]. Moreover, we demonstrated that *Tb*PARP colocalizes with these structures (Supplementary Figure 1). This is, to our knowledge, the first report of a PARP enzyme associated with the kinetoplast through the basal body.

**Figure 4.**
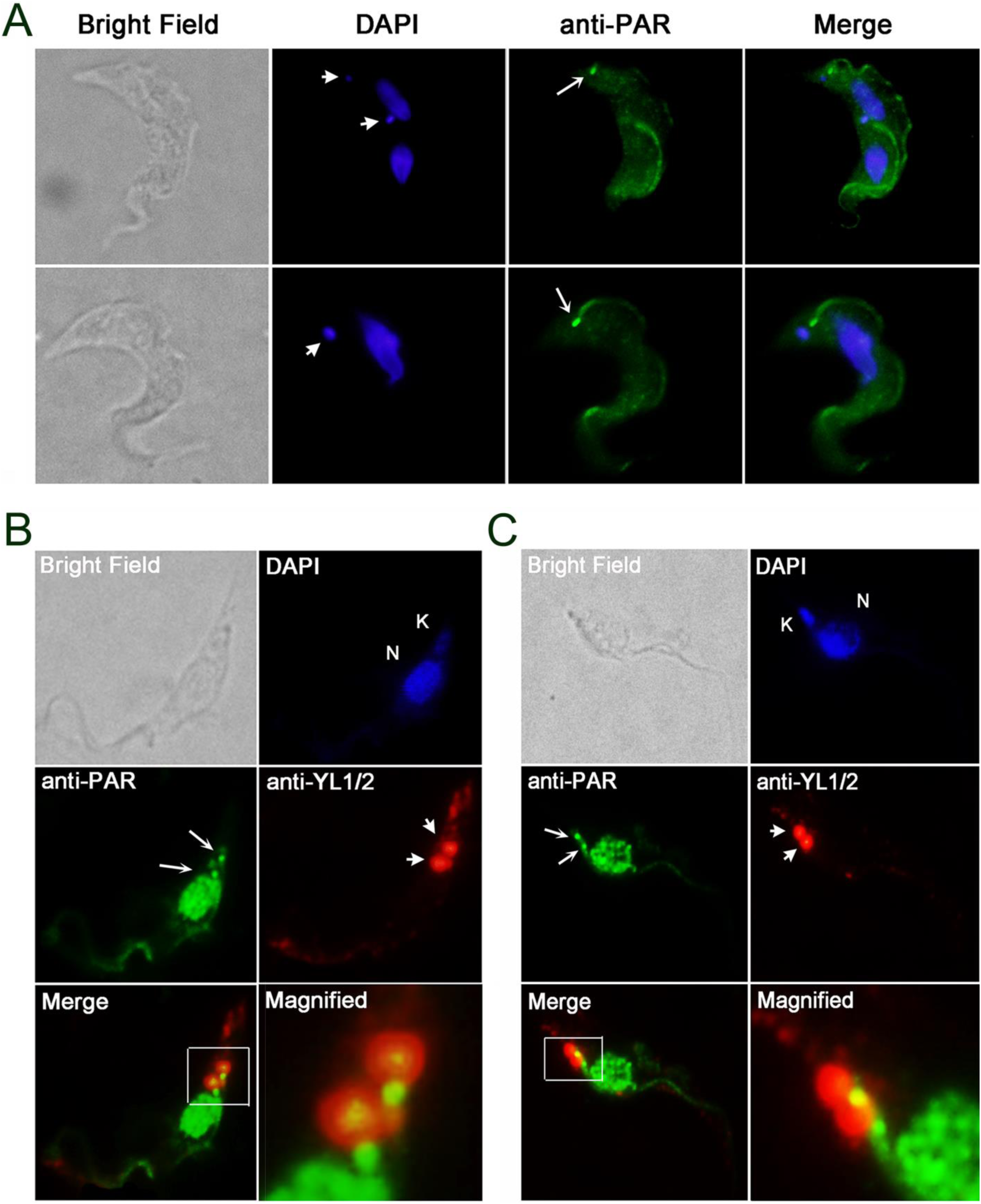
**Indirect immunofluorescence of wild type parasites.** (A) PAR was detected with specific polyclonal antibodies (arrow). DAPI was used to identify nuclear and kinetoplastic DNA (arrowhead). (B) Cytoskeleton structures from parasites were fixed with methanol and marked with anti-PAR antibody (arrow). The mature basal body was detected with anti-YL1/2 antibody (arrowhead). DAPI was used to identify nuclear (N) and kinetoplastic (K) DNA. The number of cells counted per field was about 9-12 in at least two independent experiments.

The kinetoplast is physically connected to the mature basal body and pro-basal body by a tripartite attachment complex, which comprises several filaments that guide kinetoplast segregation during mitosis [8]. It is known that inhibition of basal body duplication or segregation arrests completely the cell cycle. In order to discard a potential anomalous localization of PAR in the cytoplasm of p2216-PARP lines and, therefore, a possible disruption of kinetoplast segregation; synchronized cultures were assessed by IFI with specific antibodies along the cell cycle (Fig 5). As it was evidenced in previous experiments, the polymer signal was specially observed at a spot nearby the kinetoplast in induced and control lines. Moreover, kinetoplast division did not seem to be altered in either culture. On the other hand, nuclear PAR was detected at all the times analyzed in synchronized induced cultures, but not in uninduced parasites (Fig 5).

**Figure 5.**
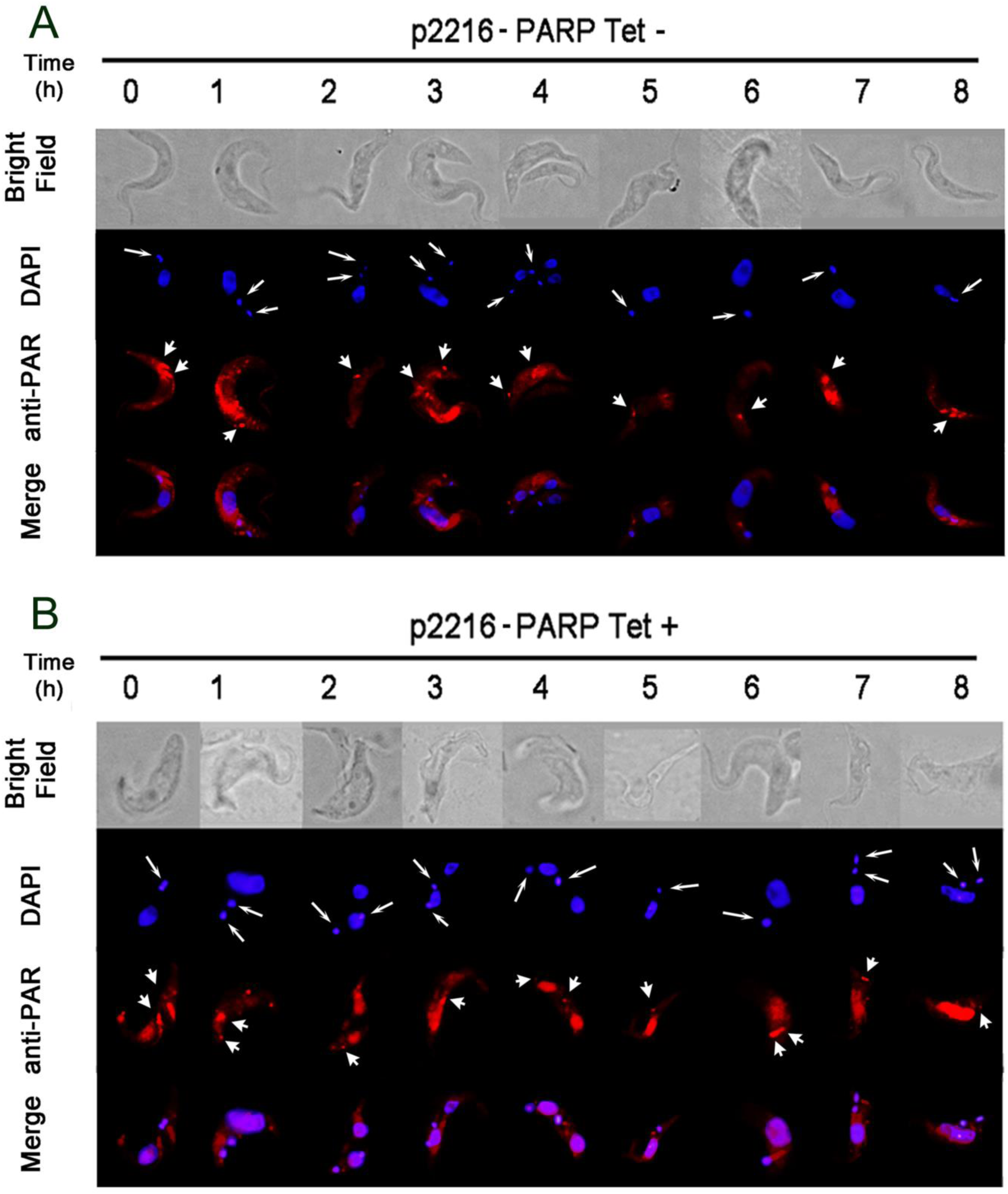
**Indirect Immunofluorescence of p2216-PARP cultures synchronized with hydroxyurea (HU).** Indirect Immunofluorescence of three day-induced (Tet +) and uninduced (Tet-) TbPARP over-expressing (p2216-PARP) cultures synchronized with 0.2 mM hydroxyurea (HU). After HU removing, parasite samples were taken every hour. Subcellular localization of the polymer was detected with anti-PAR polyclonal antibody, and the cytoplasmic signal (arrowhead) close to the kinetoplast is specifically indicated. DAPI was used to identify nuclear and kinetoplastic DNA (arrow). Representative of at least two independent experiments.

Even in absence of genomic stress, we have demonstrated that the over-expressed TbPARP-eYFP fusion protein localizes in the nucleus in p2216-PARP transgenic parasites [2]. Nuclear TbPARP-eYFP occurrence along the cell cycle in synchronized three day-induced cultures was followed by the eYFP fluorescence, presenting an evident signal in the nucleus at every time point analyzed (Supplementary Figure 2). This is in concordance with the nuclear polymer accumulation observed in the previous experiment. Fluorescent regions were also detected in the cytoplasm of both induced and control parasites. In the latter case the endogenous *Tb*PARP was identified by IFI using specific antibodies (Supplementary Figure 2). Something similar was observed in synchronized RNAi-PARG transgenic lines. While the cytoplasmic localization pattern of PAR remained unmodified in the induced and uninduced cultures, a strong polymer signal was found in the nucleus of the three-day induced parasites in every phase of the cell cycle (Fig 6). The use of HU could be the reason why PAR was produced in this organelle by the endogenous *Tb*PARP, since this compound would stimulate its function by damaging the DNA. Down-regulation of *Tb*PARG expression would avoid the polymer hydrolysis, resulting in a PAR nuclear accumulation. In contrast, control parasites that were not deprived of *Tb*PARG did not present a nuclear polymer signal.

**Figure 6.**
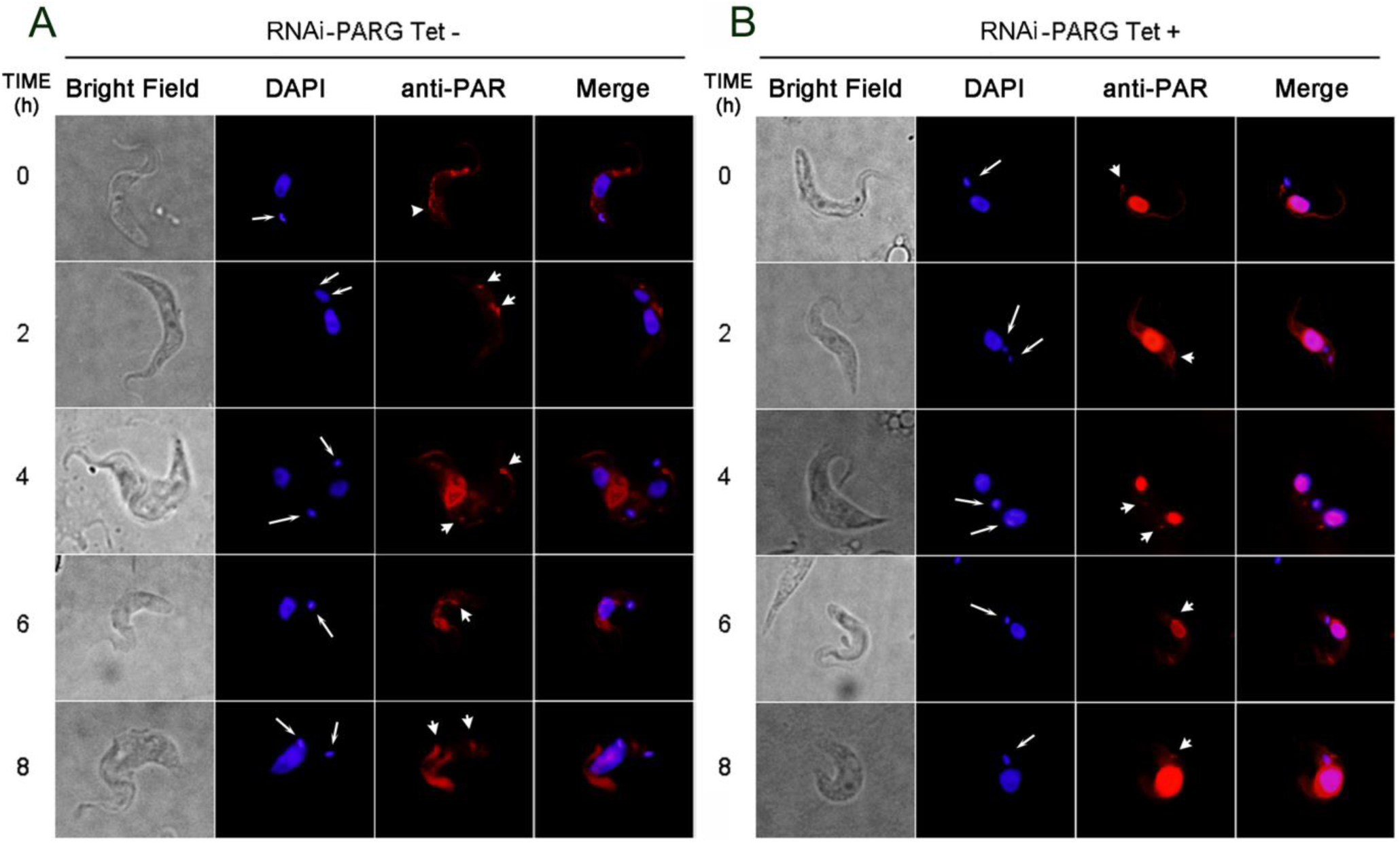
**Indirect Immunofluorescence of RNAi-PARG cultures synchronized with hydroxyurea (HU).** Indirect Immunofluorescence of three day-induced (Tet +) and uninduced (Tet-) TbPARG down-regulated (RNAi-PARG) cultures synchronized with 0.2 mM hydroxyurea (HU). After HU removing, parasite samples were taken every two hours. Subcellular localization of the polymer was detected with anti-PAR polyclonal antibody, and the cytoplasmic signal (arrowhead) close to the kinetoplast is specifically indicated. DAPI was used to identify nuclear and kinetoplastic DNA (arrow). Representative of at least two independent experiments.

Altogether, the results obtained in this work showed that p2216-PARP and RNAi-PARG transgenic lines have PAR accumulated in the nucleus in every phase of the cell cycle. However, only the passage through the G1 phase seemed to be affected by the presence of the polymer in this organelle. This delay in the cell cycle progression would explain the diminished growth rate of the transgenic trypanosomatid cultures. Finally, we showed for the first time that PAR metabolism could be related to the kinetoplast through the basal body. Further studies should be carried out in order to unravel the nexus between PAR metabolism and the kinetoplast.

## 4 Discussion

The transgenic line p2216-PARP of procyclic *T. brucei* cultures, previously obtained in our laboratory, over-express *Tb*PARP-eYFP. This fusion protein migrates always to the nucleus, where it synthesizes PAR. The same response is triggered by the genotoxic compound hydrogen peroxide in wild type parasites and at high concentrations of this agent, death takes place through an apoptotic-like pathway. The transgenic cultures, on the other hand, died through a necrotic mechanism when exposed to the same concentration of hydrogen peroxide. This observation indicates that, in this trypanosomatid, nuclear PAR may be acting as a molecular switch between different cell death pathways.

*T. brucei* RNAi-PARG transgenic cultures present a very similar phenotype to p2216-PARP parasites. This is not surprising since the absence of a functional *Tb*PARG protein also leads to PAR accumulation in the nucleus. The ratio of live and dead parasites was similar in both transgenic lines compared to the uninduced control groups. Their morphology assessed by fluorescence microscopy did not vary either. Even though we have studied the effect of an excess of nuclear polymer in procyclic parasites under conditions of genomic stress, we have not analyzed yet the consequences of a modified PAR phenotype in parasites under standard conditions.

In the present work we confirmed by a motility assay and by flow cytometry with PI and AnnexinV-FITC stains that PAR-altered levels did not cause an increase in the rate of parasite cell death. However, we observed that a modification in the PAR balance produced a differential growth rate in procyclic p2216-PARP and RNAi PARG cultures. Indeed, the number of the control (uninduced) cells was almost twice as much as the number of the induced parasites at day three. Further studies demonstrated that this retardation in the growth curve obeyed to a slowed down cell cycle.

Cell cycle in procyclic *T. brucei* parasites is unique in many aspects. At the end of the G_1_ phase, the kinetoplast has already duplicated and the pro-basal body has become a mature basal body. During the S phase the nuclear DNA replicates while the two kinetoplasts start separating from each other with their pro-basal and basal bodies. The formation of a novel flagellum is initiated in this phase as well. At the beginning of the G2/M phase the kinetoplasts are visibly distant and the nuclear division takes place after the mitotic spindle emergence. The flagellum continues elongating until the cytokinesis, produced at the end of the G2/M phase.

Synchronized cultures were analyzed by flow cytometry and Propidium Iodide stain. Procyclic wild type parasites are reported to arrest at the end of the S phase when incubated with HU. This was actually what we have observed with the uninduced control groups of the p2216-PARP and RNAi-PARG transgenic lines. However, in the induced cultures we evidenced a largerG1 population immediately HU removing (time 0). One possible explanation is that incubation in the presence of HU could cause death of the induced parasites after being stalled in G1 phase at 0 hour. We have previously seen that another genotoxic agent, hydrogen peroxide, renders PAR-accumulated parasites more sensitive to DNA stress. Another explanation could be that the induced cultures progress through the cell cycle so slowly that the synchronization time is not enough to arrest them at the S phase.

Later on, after HU was moved the induced parasites present a larger G1 population respect to the uninduced counterparts at certain hours, which indicates that they are impeded to progress beyond the G1 stage.

We did not evidence any difference of PAR and *Tb*PARP cytoplasmic signals between synchronized induced parasites and control groups along the cell cycle; similarly, kinetoplast migration did not seem to be altered. It is interesting that *Tb*PARP colocalizes with the mature basal body and that PAR occurs next to this structure. The basal body is a centriole-like structure which is situated in the base of the flagellum. The kinetoplast is physically connected to the proximal end of both the mature basal body and pro-basal body by a tripartite attachment complex [8]. Such a complex comprises several filaments that guide kinetoplast segregation during mitosis [9]. It has been published that even though problems in the mitotic phase do not halt cytokinesis, inhibition of basal body duplication or segregation arrests completely the cell cycle at this stage. Indeed, it was proposed that basal body migration could function as the main driving force of cytokinesis. We have not observed that cell cycle progression was arrested at the G2/M phase in this work, which is in line with the evidence that basal body and kinetoplast migration did not seem to be disturbed in the transgenic lines p2216-PARP and RNAi-PARG.

On the other hand, sequence analysis determined that *Tb*PARP presents a high grade of similarity with the human PARP-3 [1].This protein has been identified as a central component of the centrosome, and it is preferably located in the new centriole along the human cell cycle [9] Even more, it has been reported that hPARP-3 influences negatively cell cycle progression in the G1/S boundary, without interfering with centrosome duplication [10]. How nuclear PAR could control the G1/S transition in procyclic *T. brucei* is an open question that should be addressed in the future.

Despite the fact that nuclear polymer signal is detected in induced cultures in every phase of the cell cycle, PAR accumulation in the nucleus only affects them at the G1 stage. Parasites have trouble to progress through the G1/S transition. *T. brucei* possesses cyclins and cyclin-dependent kinases generically termed CDC2-related kinases (CRKs) [11]. Some of them are key regulators of the cell cycle [12] and play a role in G1/S and G2/M transitions, with no homologs found in humans [13, 14]. It would be interesting to identify in the future whether some of these proteins interact with *Tb*PARP or are target of poly(ADP-ribosyl)ation. Further studies are needed in order to elucidate the role that PAR plays within the cell cycle, especially at the stage G1.

## Acknowledgements

This work was supported by Agencia Nacional de Promoción Científica y Tecnológica (Argentina), Consejo Nacional de Investigaciones Científicas y Técnicas (Argentina) and Universidad de Buenos Aires (Argentina).

